# Reconstitution of the host holobiont in germ-free rats acutely increases bone growth and affects marrow cellular content

**DOI:** 10.1101/2020.07.15.201657

**Authors:** PJ Czernik, RM Golonka, S Chakraborty, BS Yeoh, A Abokor, P Saha, JY Yeo, B Mell, X Cheng, S Baroi, B Joe, M Vijay-Kumar, B Lecka-Czernik

## Abstract

In recent years there has been growing evidence regarding the effect of microbiota on the skeletal growth and homeostasis. Here we present, for the first time, accelerated longitudinal and radial bone growth in young (7-week-old) germ-free male rats after short-term exposure to a newly established gut microbiota. Changes in bone mass and structure were analyzed after 10 days following the onset of colonization through cohousing with conventional rats and revealed unprecedented acceleration of bone accrual in cortical and trabecular compartments, increased bone tissue mineral density, improved proliferation and hypertrophy of growth plate chondrocytes, bone lengthening, and preferential deposition of periosteal bone in tibia diaphysis. In addition, the number of small-in-size adipocytes increased, while the number of megakaryocytes decreased, in the bone marrow of conventionalized germ-free rats. The observed changes in bone status were paralleled with a positive shift in microbiota composition towards short chain fatty acids (SCFA)-producing microbes, which reflected a dramatic increase in cecal concentration of SCFA, specifically butyrate. Further, reconstitution of the host holobiont increased hepatic expression of IGF-1 and its circulating levels, implicating an involvement of the somatotropic axis. Increased serum levels of 25-hydroxy vitamin D and alkaline phosphatase pointed toward an active process of bone formation. The acute stimulatory effect on bone growth occurred independently of body mass increase and resembled reversal of dysbiosis in adolescence, which is marked by rapid skeletal expansion. These findings may help in developing microbiota-based therapeutics to combat bone related disorders resulting from hormonal defects and/or malnutrition in children and adolescence.

## Introduction

Coined by Lynn Margulis in 1991, the term holobiont describes how macro-species, such as mammals, live in symbiosis with micro-species, whereby all participating individuals are bionts and the entire organism comprised of these bionts is a holobiont (1,2). An animal is therefore a holobiont by hosting a diverse set of gut microbes whose bacterial genome outweighs the host genome by 150-fold, but the bacteria and host cell number ratio is recently calculated to be 1:1 compared to previous notions (3). Commensal microbes have co-evolved with the host and thus, have become a vital component of a comprehensive ecosystem that orchestrates normal physiology of the holobiont. Consequently, an alteration in the composition of the micro-species can negatively influence host physiology, which has prompted the concept of reconstituting or correcting the gut microbiome as a therapeutic agent (e.g. probiotics). Extensive studies in animal models and ongoing clinical trials involving the microbiota (4) provide promising means of medical intervention to mitigate incurable conditions and augment existing therapies for the treatment of recurring intestinal infections, bowel diseases, obesity, type 2 diabetes, osteoporosis, and osteoarthritis (5-8).

There is a general consensus that gut microbiota influence bone biology; however, distinct pathways connecting these two entities make the nature of this interaction diverse and complex. Intestinal microenvironments comprised of host- and bacterial-derived metabolites and viable bacteria, which later regulate the gut-immune network, could lead to seemingly various and contrasting observations in bone phenotype. For instance, butyrate, which is a colonic metabolite from microbial fermentation of complex fermentable carbohydrates, signals to bone in discrete modes. Butyrate can stimulate bone formation by an expansion of intestinal- and bone marrow-derived regulatory T cells (T_reg_), where the interaction with marrow CD8^+^T cells results in increased expression of the bone anabolic WNT10b ligand (9). Simultaneously, butyrate can act on the somatotropic axis by increasing growth hormone release from the pituitary, leading to increased presence of circulating insulin-like growth factor-1 (IGF-1) (10), which may drive the expansion of body and bone mass independently or in concert with the effect of butyrate on WNT signaling.

Experimental conditions, like the animal model, strain, sex, age, housing environment (including the diet), and even the animal source can influence outcomes of studies focused on the gut microbiota-bone axes (11). Importantly, postnatal development studies using young mice born in germ-free (GF) conditions show that the dynamics of bone growth can be differently affected by the animals genetic background, age, and route of microbiota delivery. For example, a study by Yan et al. shows that gut colonization of 8-week old, germ-free BALB/c and C57BL/6 hybrid mice with feces collected from their non-GF genetic counterpart led to initial loss of bone mass followed by a slow gain over the next 8 months (12). However, Sjögren et al. show relatively faster gain of bone mass in 3-week old, germ-free C57BL/6 mice after colonization with fresh cecal transfer derived from their non-GF genetic counterpart (13). In further contrast, intragastric gavage of microbiota derived from human, mouse, or rat to 4-week old mice did not have any effect on bone mass (11,14-16). One of the possible explanations of such a variety of effects may include recipient response to the foreign microbiome. Direct transfer of established microbiota, which carries a physiologic imprint of its original host, may lead to adverse effects in GF animals as it is recognized as ‘infection’, and results in immune response and inflammation. This offensive state persists to the point when a new host and microbiota adapt to each other. In the model presented here, cohousing of GF with conventionalized rats allows for the gut to be gradually populated by microbes and avoids a bolus effect associated with direct microbiota transplantation.

An important question remains unanswered, which is does gut microbiota participate in host bone growth either as a ‘passive’ commensal entity or rather an ‘active’ organ that constitutes an integral physiologic component of the body to support bone metabolism and homeostasis. In the presented study, we aimed to explore this gap in the field by answering the following questions: 1) can a GF rat model be used to study the effect of gut microbiota on bone growth?; 2) is the transfer of microbiota through cohabitation of GF rats with non-GF rats advantageous on bone growth in respect to natural gut colonization via coprophagy?; and 3) is this model applicable to human conditions of stunted growth? We compared 7-week old rats born and raised in GF conditions with the sex- and age-matched counterpart exposed to gut microbiota by co-housing with holobiont intact rats over a short, 10-day time period. Our study shows that the gut microbiome is essential for skeletal maturation in an adolescent organism, as the restoration of micro-species led to an immediate resumption of bone growth that had been stunted in gut microbiota depleted GF animals. In conclusion, the model of gut microbiota colonization in GF rats by cohabitation with holobiont intact rats is robust and can be very useful to study stunted bone growth found in juveniles with microbiota dysbiosis.

## Materials and Methods

### Animals

All animal experiments were conducted according to the University of Toledo Institutional Animal Care and Use Committee approved protocols. This study was conducted using male, 7-week-old Sprague Dawley (SD) rats that were concomitantly raised as germ-free rats until the commencement of the study. Germ free rats were separated into two groups, either germ-free (GF) (n=6/group) or germ-free conventionalized (GFC) (n=6/group). These animals were raised and set up for studies at Taconic Biosciences (Rensselaer, NY). Conventionalization of GFC rats were performed by co-housing GF rats with conventionally-raised (C) SD rats for 10 days (1:1 ratio). Upon arrival at the University of Toledo, the animals were immediately used for experiments. The animals were euthanized via CO_2_ asphyxiation, blood and tissues were collected and stored at −80°C until further use.

### Assessment of microbiota composition

The analysis of the microbiota composition and biodiversity in C, GF, and GFC animals was done by 16S ribosomal RNA gene as previously described (17), and as reported for exactly the same set of animals in a concurrent recently published paper (18). Tissue collection for experiments described in the above publication and for this manuscript was conducted at the same time in a joint effort of several laboratories, and coordinated by the Microbiome Consortium at the University of Toledo.

### Collection and processing of specimens

Femurs and tibias extracted from euthanized animals were thoroughly cleaned of muscle tissue. Both tibias from each animal were immediately preserved in 10% NBF for subsequent evaluation of bone microstructure and for preparation of histological sections. For RNA isolations from bone and marrow, single femurs were kept on ice after extraction and processed as follows. Proximal and distal ends were cut off using rotary diamond micro saw, diaphysis bone sections were suspended in a hole drilled in microtube lid, and then bone and marrow were separated by centrifugation at 2000 x g for 1 min at 4°C. Following separation, bone and marrow were flash frozen in liquid nitrogen. Tibias intended for preparation of histological sections were kept in 10% NBF for 3 days, then decalcified on a rocking platform with Formical-4 reagent (StatLab, McKinney, TX, USA) in 3 consecutive extractions lasting 24, 6, and 2 hours using 20 ml of the decalcifier. Oxalic acid test was conducted on the last fraction to ensure the complete calcium removal and decalcified bones were stored in 10% NBF for further processing. Cecal short chain fatty acids (SCFA) levels were measured by 1H-NMR, as described (19).

### Bone morphometry

Assessment of trabecular bone in proximal tibia and cortical bone at tibia midshaft was conducted by micro CT using μCT 35 system (Scanco Medical AG, Bruettisellen, Switzerland). Bone scans were performed with the x-ray source operating at 70 kVp and 114 μA energy settings, and recording 500 projections/180°acquired at 300 ms integration time using 20 μm nominal resolution voxel for both bone locations. Scans of the proximal tibia consisted of 340 cross-sectional slices starting at the top of growth plate, and images of trabecular bone were segmented at optimized lower threshold value of 220 using per mille scale (approximately 3000 HU, or μ of 1.76) following manual contouring starting 10 slices down from the intercondylar notch and extending for 200 more slices. Scans of the cortical bone at tibia midshaft contained 57 slices all of which were contoured automatically, and which were segmented at 260 per mille threshold. Computed bone bending strength (I_max_/C_max_ and I_min_/C_min_) and torsional strength (pMOI) at tibia midshaft were based on bone cross-sectional geometry in combination with local tissue mineral content. The analysis of the trabecular bone microstructure, cortical bone parameters, and bone strength, was conducted using Evaluation Program V6.5-3 (Scanco Medical AG) and conformed to recommended guidelines (20).

### Gene expression

Total RNAs from bone tissue and marrow specimens were isolated using TRI Reagent (MRC Inc., Cincinnati, OH, USA) following manufacturers’ protocol. Frozen marrow and bone were directly homogenized in TRI Reagent using microtube pistons and rotor-stator homogenizer, respectively. cDNAs were synthesized using 0.25 μg of isolated RNAs and Verso cDNA Synthesis Kit (Thermo Fisher Scientific, Waltham, MA, USA). Oligonucleotide primers for real-time qPCR were designed using Primer-Blast (NCBI, NIH) with PCR product length set from 80 to 120 nucleotides and 60°C optimal primer melting temperature. Oligonucleotides were synthesized by the Integrated DNA Technologies, Inc. (Coralville, IA, USA) and melting temperatures, adjusted for qPCR conditions, were calculated using OligoAnalyzer Tool available at the manufacturer’s website. Oligonucleotide sequences, amplicon sizes, and melting temperatures are listed in the Supplementary Table 1. qPCR amplifications were carried out using TruAmp qPCR SYBR Green SuperMix-Rox (Smart Bioscience Inc., Maumee, OH, USA) and StepOne Plus system (Applied Biosystems Inc., Foster City, CA, USA). Amplifications were carried out using 40 cycle two-step amplification protocol with annealing temperatures set to 4-5 degrees below calculated primers melting temperatures, and finished with melting curve cycle to verify homogeneity of amplification products. Relative gene expression was analyzed by the comparative ΔΔCT method using *18S* rRNA levels for normalization.

### Histology and image quantification

Decalcified tibia bones were segmented at tibia crest and tibiofibular junction, and the resulting proximal and mid diaphysis fragments were embedded in paraffin blocks and processed at the University of Toledo Histology Core Facility. High resolution images of longitudinal 6 μm sections stained with H&E were generated using Olympus VS120 slide scanner. Images in the native VSI format were converted to TIFF images at the resolution of 0.685 μm/pixel which were then used for histological analysis. Image conversion and subsequent measurements of circumferential lamellar bone, growth plate, and marrow adiposity were conducted using tools available in Fiji-ImageJ image processing package (21) equipped with OlympusViewer Plugin (Olympus Corp., Tokyo, Japan). Determination of adipocyte count and cell size distribution in bone marrow was done on 8-bit grayscale images after applying grayscale threshold of 210-254 which effectively separated adipocytes from the majority of bone marrow cellular components (histogram peak). Since bone marrow adipocytes significantly differ in size, 3 random bone marrow 0.3 mm^2^ areas were sampled and it was determined that histologically relevant single adipocyte area was within the range of 90-1400 μm^2^. In order to eliminate irregular objects of similar areas roundness threshold was set to 0.4, which was equivalent to a 2.5 aspect ratio. Megakaryocyte number was assessed by direct counting over entire histological sections of mid diaphysis stained with H&E, and the cells were distinguished from other cellular components of the marrow by size and the presence of multilobated nuclei.

### Serum chemistry

C, GF and GFC rats were euthanized by CO_2_ asphyxiation and blood was collected *via* cardiac puncture in BD microtainer (Becton, Dickinson, Franklin Lakes, NJ). Hemolysis-free sera were obtained after centrifugation 10,000 rpm, 10 min, 4°C and stored at −80°C until further use. Automated assessment of serum markers was conducted using VetScan 2 analyzer (Abaxis, Inc., Union City, CA, USA) and Comprehensive Diagnostic Profile rotor (Abaxis 500-7123) which contains tests for alanine aminotransferase (ALT), albumin (ALB), alkaline phosphatase (ALP), amylase (AMY), total calcium (Ca), creatinine (CRE), globulin (GLOB), glucose (GLU), phosphorus (Phos), potassium (K), sodium (Na), total bilirubin (TBIL), total protein (TP), and blood urea nitrogen (BUN). Serum preparation and measurements were conducted according to a protocol provided by the manufacturer. Serum IGF-1 levels were quantified by an ELISA kit from R & D Systems (Minneapolis, MN) according to kit instructions.

### Statistical analysis

Data are presented as means + SD. Statistical analysis was performed using two-tailed Student’s test and Pearson correlation to compare animal groups using GraphPad Prism 8.3 (GraphPad, La Jolla, CA, USA). For multiple comparison one-way Anova was performed. Statistical differences with p < 0.05 were considered significant. Statistical analysis of microbial α- and β-diversity was performed, as previously described (17).

## Results

### Metagenomics signify a distinct microbial composition after bacterial colonization in conventionalized germ-free (GFC) rats

Assessment of microbiota composition revealed successful colonization of GFC rats with some distinctive features present in conventional (C) rats. Total bacterial 16S copy number per fecal pellet was the same in C and GFC animals at 4.5×10^8^, while in GF animals the detected copy number was 30,000-fold lower at 1.5×10^4^. There was no significant difference in α-diversity between C and GFC animals, meaning that the number of different species was comparable in both groups (18). However, significance in β-diversity was observed in compositions of microbial communities between C and GFC animals, including low-abundance taxa (Table 1). Remarkably, relative abundance of *Clostridiaceae* and *Clostridium* taxa was significantly increased in GFC versus C animals by 3.2- and 13.4-fold, respectively, with higher taxa (Firmicutes, Clostridia, and Clostridiales) significantly decreased in GFC versus C animals by 2-fold. Importantly, the relative abundance of *Clostridium* spp. in GFC rats was comparable with the abundance of Clostridium_SMB53 observed in normal NIH heterogenous stock rats (22). This observation indicates a specific compositional shift toward *Clostridium* spp., which occurred during colonization of the GF animal with C rat microbiota. A modest, but statistically significant, increase in relative abundance was observed for the *Muribaculaceae* family (formerly known as S24.7), which accounted for the 100% increase in abundance of Bacteroidetes phylum. Moreover, a 4-fold increase in relative abundance of the Proteobacteria phylum in GFC animals was exclusively represented by expansion of *Suterella* spp. (Table 1).

**Table 1.** Comparison of gut microbial composition of C and GFC rats

| Phylum | Class | Order | Family | Genus | Relative abundance |  |  | Fold change |  |
| --- | --- | --- | --- | --- | --- | --- | --- | --- | --- |
|  |  |  |  |  | C | GFC | p value | Up | Down |
| Firmicutes |  |  |  |  | 0.5727 | 0.3570 | 0.025 |  | 1.6 |
| Firmicutes | Clostridia |  |  |  | 0.5442 | 0.3058 | 0.010 |  | 1.8 |
| Firmicutes | Clostridia | Clostridiales |  |  | 0.3582 | 0.1223 | 0.007 |  | 2.9 |
| Firmicutes | Clostridia | Clostridiales |  |  | 0.5440 | 0.3055 | 0.010 |  | 1.8 |
| Firmicutes | Clostridia | Clostridiales | Clostridiaceae |  | 0.0135 | 0.0432 | 0.040 | 3.2 |  |
| Firmicutes | Clostridia | Clostridiales | Clostridiaceae | Clostridium | 0.0021 | 0.0281 | 0.007 | 13.4 |  |
| Proteobacteria |  |  |  |  | 0.0279 | 0.0973 | 0.060 | 3.5 |  |
| Proteobacteria | Betaproteobacteria |  |  |  | 0.0227 | 0.0938 | 0.040 | 4.1 |  |
| Proteobacteria | Betaproteobacteria | Burkholderiales |  |  | 0.0227 | 0.0938 | 0.040 | 4.1 |  |
| Proteobacteria | Betaproteobacteria | Burkholderiales | Alcaligenaceae |  | 0.0227 | 0.0937 | 0.040 | 4.1 |  |
| Proteobacteria | Betaproteobacteria | Burkholderiales | Alcaligenaceae | Sutterella | 0.0227 | 0.0937 | 0.040 | 4.1 |  |
| Bacteroidetes |  |  |  |  | 0.3439 | 0.4521 | 0.060 | 1.3 |  |
| Bacteroidetes | Bacteroidia |  |  |  | 0.3439 | 0.4521 | 0.060 | 1.3 |  |
| Bacteroidetes | Bacteroidia | Bacteroidales |  |  | 0.3439 | 0.4521 | 0.060 | 1.3 |  |
| Bacteroidetes | Bacteroidia | Bacteroidales | Muribaculaceae |  | 0.3035 | 0.4272 | 0.030 | 1.4 |  |

### Elevated levels of serum markers related to bone mineralization and liver function in GFC rats

To assess systemic features after the presence of a newly established microbiota in GFC rats, we analyzed 13 blood serum markers and electrolytes indicative of bone, liver, and kidney function (Table 2). Most of the assessed markers remained at comparable levels in GF and GFC animals, and were considered normal when compared to reference values published for normal wild-type, male Sprague-Dawley rats (23-25). These included protein and electrolyte markers, namely amylase (AMY), creatinine (CRE), total protein (TP), phosphorous (Phos), sodium (Na) and potassium (K) (Table 2). Among the most prominent features was serum albumin (ALB), which was elevated in GF animals by 75%, as compared to reference values, and was significantly reduced in GFC rats to being elevated by only 48% (Table 2, Fig. 1A). There is very limited information regarding hyperalbuminemia in animals and humans except that it is a rare condition mainly associated with dehydration. However, globulin (GLOB) level, *albeit* markedly lower than the reference value, was identical in both groups with the total serum protein decrease following that of the ALB, which indicated that dehydration was unlikely to be responsible for high ALB levels in GF rats. Alanine transaminase (ALT), a marker of liver dysfunction, and blood urea nitrogen (BUN), a marker of kidney failure and increased protein degradation in liver, were not different in both GF and GFC rats (Fig. 1B-C), but they were elevated by 75% and 52%, respectively, as compared to reference values shown in Table 2. However, creatinine (CRE) readings were standard in both groups of animals suggesting normal kidney function. These observations imply that liver function, manifested by an increase in ALT and BUN, and unexpected elevation in ALB synthesis, is affected by the lack of microbiota and is not fully restored by a short-term presence of gut microbiota, except for partial normalization of ALB level. Further, there are indicators for altered bone metabolism in GFC rats. Alkaline phosphatase (ALP) level was significantly elevated by 33% (Fig. 1D); however, serum calcium concentration was significantly decreased by 6% in GFC versus GF rats most likely as a result of increased bone formation (Fig. 1E), with serum levels of 25-OH vitamin D significantly increased by 8% (Fig. 1F) suggesting a physiologic response to increased calcium demand.

**Table 2.**
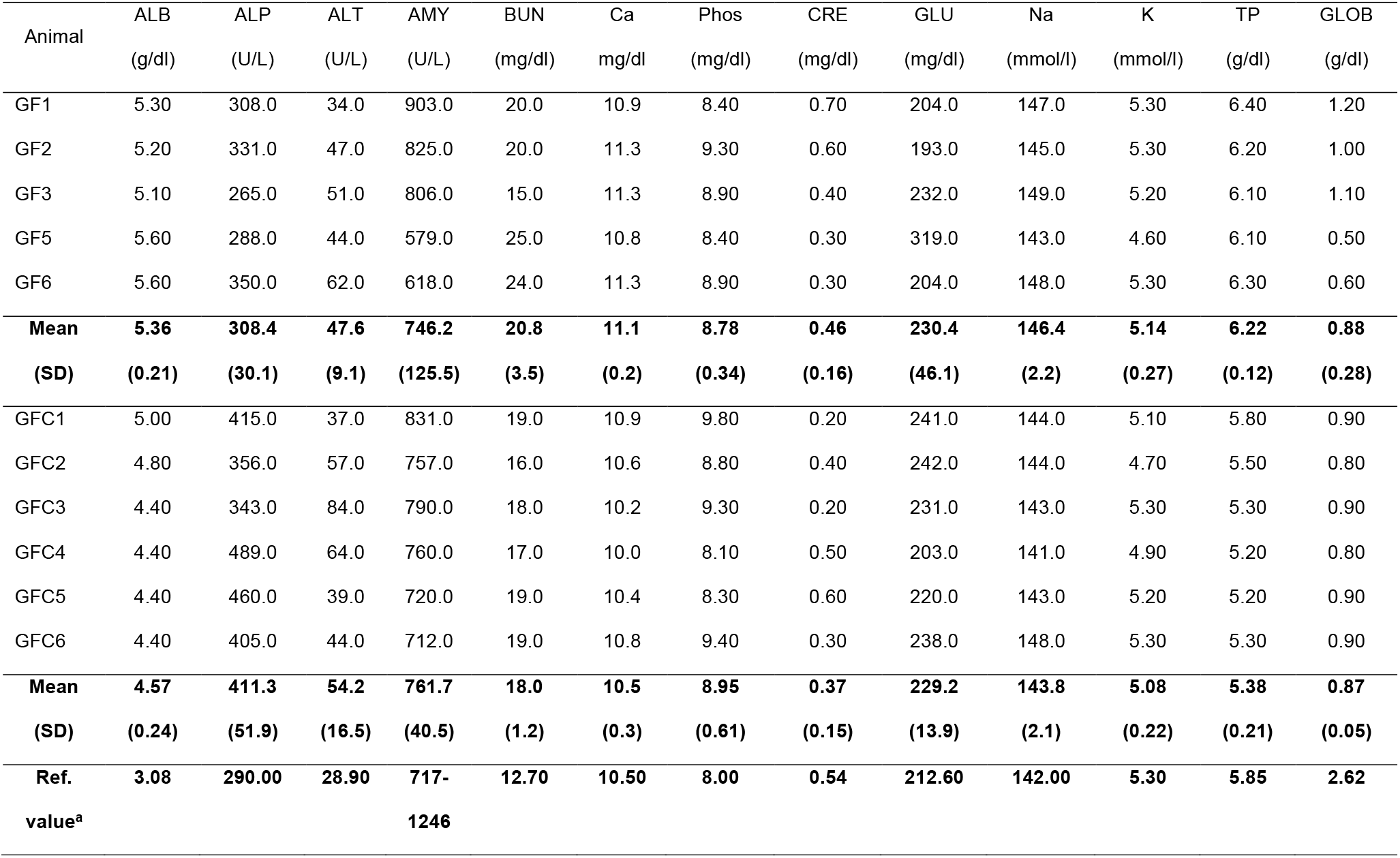
Serum comprehensive diagnostic profile. ALB – albumin, ALP - alkaline phosphatase, ALT - alanine aminotransferase, AMY – amylase, BUN - blood urea nitrogen, Ca - total calcium, Phos – phosphorus, CRE - creatinine, GLU - glucose, Na - sodium, K - potassium, TP - total protein, GLOB – globulin. SD – standard deviation. ^a)^Reference values were obtained from published sources (23-25).

**Figure 1.**
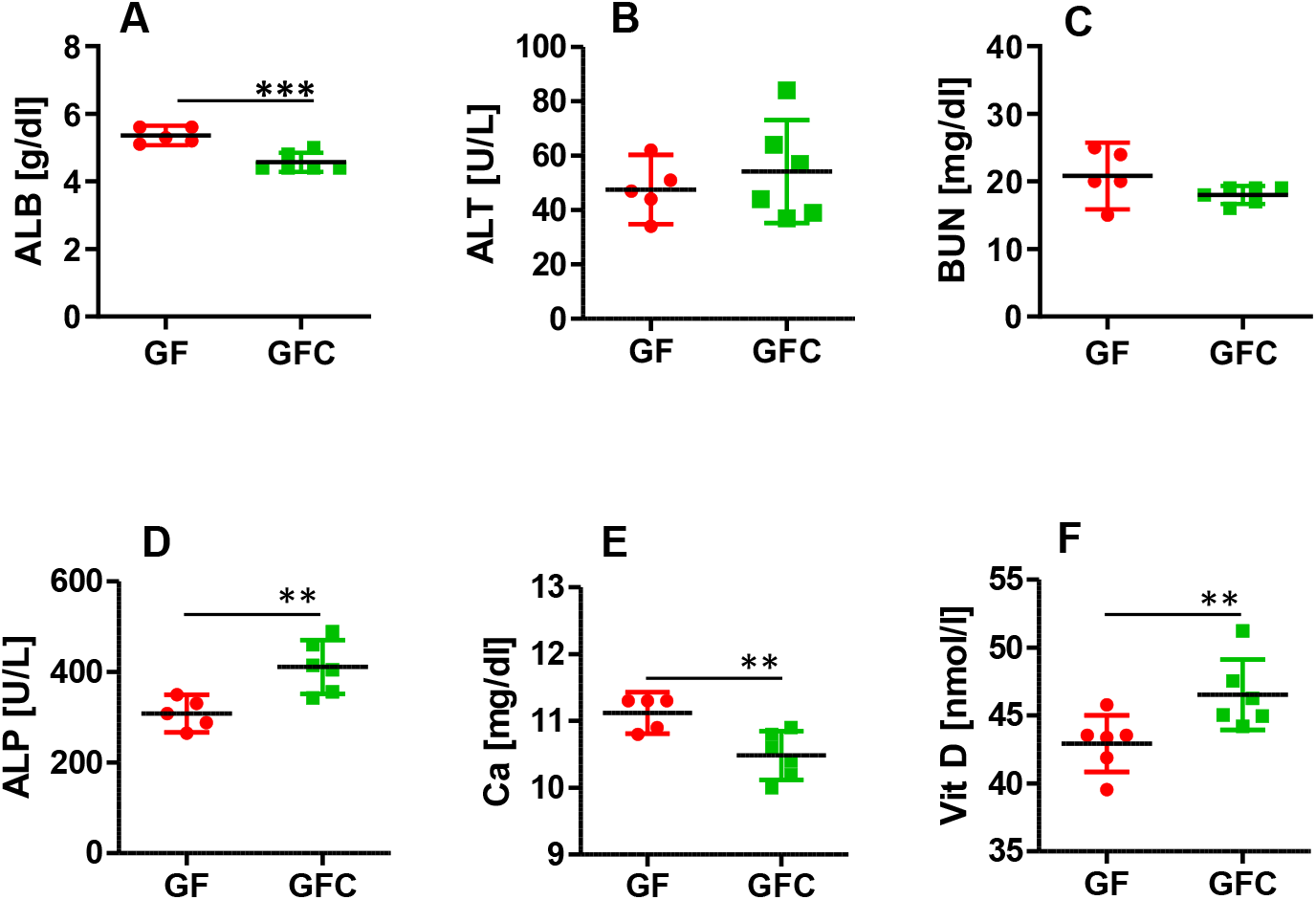
Changes in serum markers of liver function and bone mineralization in GF and GFC rats. Graphs A, B, and C respectively show liver markers concentrations of albumin (ALB), alanine transaminase (ALT), and blood urea nitrogen (BUN). Graphs D, E, and F show bone markers concentrations of alkaline phosphatase (ALP), calcium (Ca), and vitamin D. **: p<0.01, ***: p<0.001

### Reconstitution of the holobiont with microbiota increases bone mass of the host

To assess the immediate effect of gut microbiota on bone status, we evaluated trabecular bone morphometry of proximal tibia and cortical bone parameters at tibia midshaft using 3D images acquired by micro computed tomography (Fig. 2A-C). Measurements of trabecular bone in proximal tibia revealed statistical gains in total volume and bone volume with bone volume/total volume ratio remaining unchanged (Fig. 2D-F). However, there was a tendency to increased trabecular thickness and decreased trabecular spacing in GFC rats (Fig. 2G-I). These indicate that reconstitution of the holobiont resulted in overall bone expansion of tibia with proportional increase in trabecular bone compartment. Assessment of trabecular bone in L4 vertebral body showed similar trend of the bone expansion in GFC rats (Supplementary Table 2).

**Figure 2.**
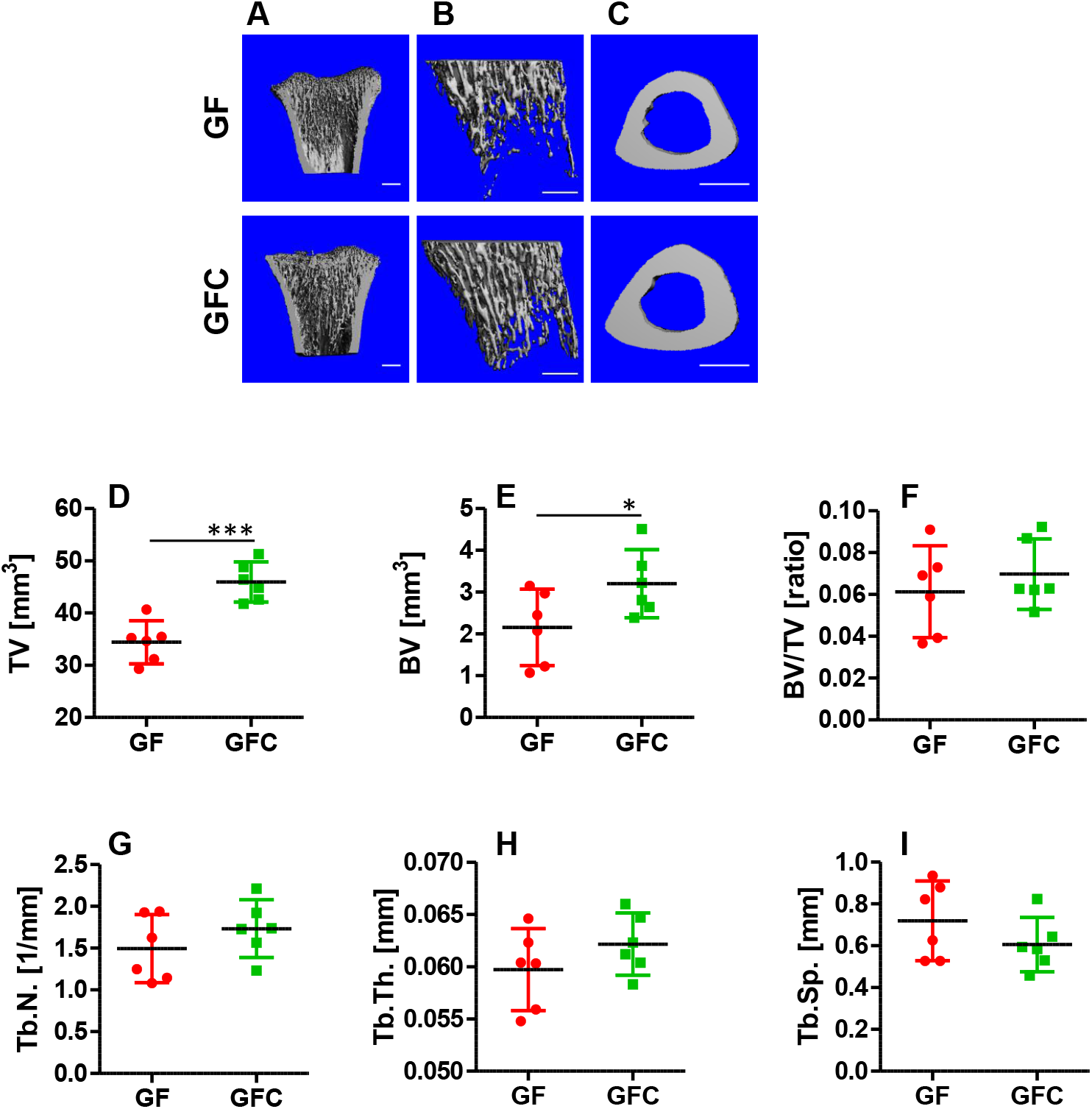
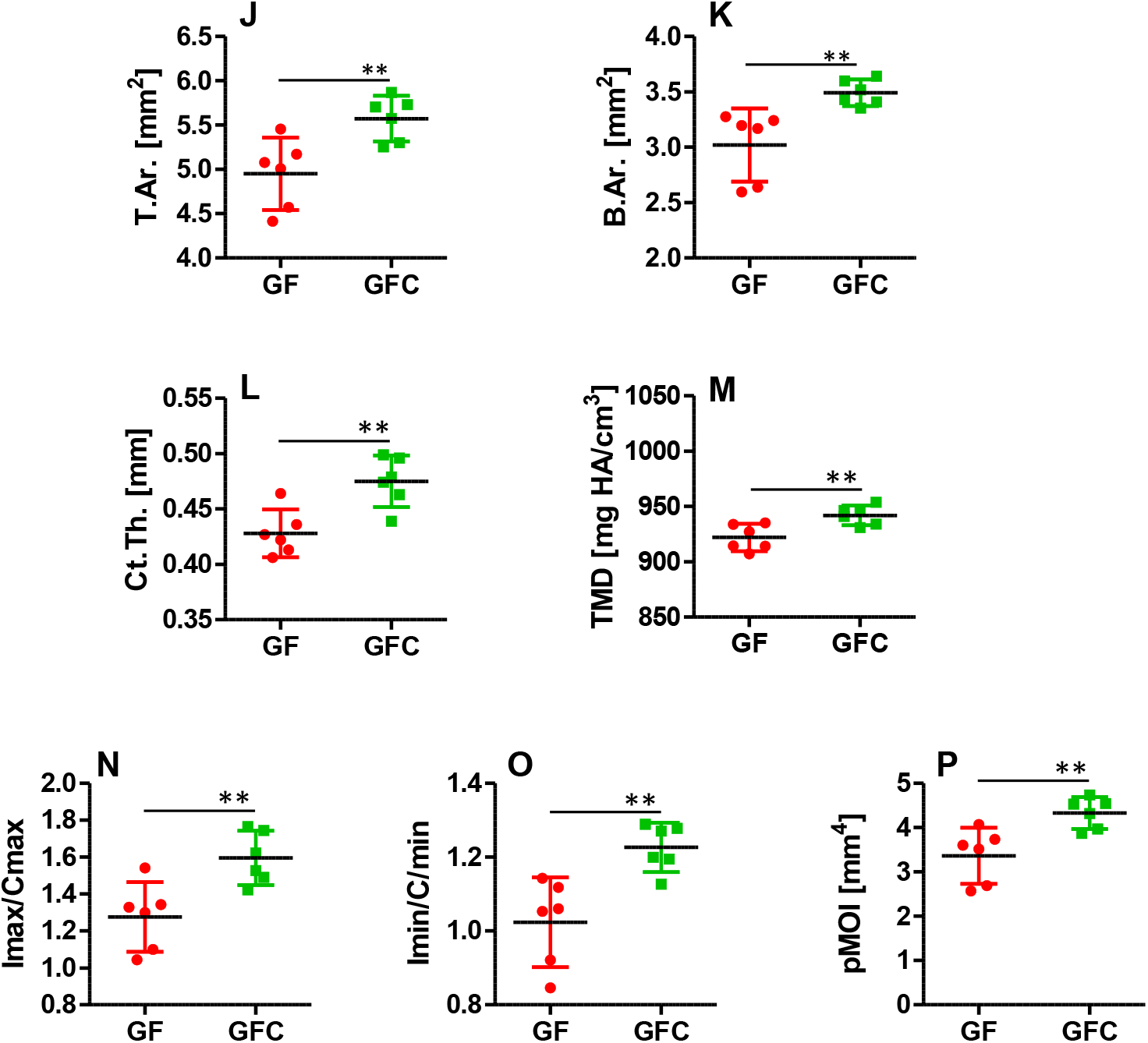
Acute effect of gut microbiota on bone mass and structure in germ-free conventionalized rats. A - C. 3D mCT bone renderings of GF rat and GFC rat show proximal tibia longitudinal sections (A), trabecular bone compartment (B), and bone cross sections at tibia midshaft (C). D - I. mCT analysis of proximal tibia trabecular bone morphometry. TV – tissue volume; BV – bone volume; BV/TV – bone fraction; Tb.N – trabecular number; Tb.Th – trabecular thickness; and Tb.Sp – trabecular spacing. J – L. Cortical bone analysis at tibia midshaft. T.Ar – tissue area; B.Ar – bone area; and Ct.Th - cortical thickness. M. Measurement of status of bone tissue mineralization. TMD – tissue mineral density. N - P. mCT analysis of tibia bone biomechanical properties. Imax/Cmax (N) and Imin/Cmin (O) are measures of predicted resistance to bending and pMOI (P) represent computed moment of inertia to torsional strength. *: p<0.05, **: p<0.01, ***: p<0.001

Cortical bone measurements revealed that within the period of 10 days, GFC rats significantly gained bone mass. Total cross-sectional area (T.Ar), bone area (B.Ar), and cortical thickness (Ct.Th) increased by 12.6%, 15.6%, and 11%, respectively (Fig. 2J-L). Marrow area (M.Ar) remained the same in both groups indicating that bone apposition in GFC rats occurred mainly on the periosteal surface (Supplementary Table 3). The average body weight for GFC rats was significantly increased by 27.3% (GF: 205.7 g + SD 30.2, GFC: 261.8 g + SD 13.3, p=0.002) suggesting that bone expansion may result from increased mechanical loading. However, while calculation of Pearson correlation coefficient (r) for cortical parameters versus body weight of GF animals showed that B.Ar (r=0.86), T.Ar (r=0.99), and Ct.Th (r=0.68) positively correlated with animal weights, in GFC animals T.Ar correlation was significantly lower (r=0.45) and B.Ar and Ct.Th correlated negatively with respective r values of -0.39 and -0.78. These data suggest that mechanical loading is not a major factor in driving cortical bone accrual. Instead, it may be associated with increased calories provided by gut microbiota fermentation of complex carbohydrates, but more importantly resulting from a molecular signaling of fermentation metabolites (e.g. SCFA) along the gut-bone axis.

It is important to stress that bone mass gain judged by an increase of mean cortical thickness in GFC rats was substantial considering the short time exposure to gut microbiota. Reported periosteal bone apposition rate in male rats of comparable age and body weight is on average 0.05 μm/h (26,27). Mean difference in cortical thickness between GF and GFC rats at the time of sacrifice was 47 μm, and gained over 10 days, giving an apparent apposition rate of 0.19 μm/h. At the same time, tissue mineral density (TMD) (Fig. 2M) in cortical bone was statistically increased in GFC rats but only by 2.1% which seemed to be less than expected when compared to the 4-fold increase in the apposition rate. Finally, calculations of bone moment of inertia at tibia midshaft showed that predicted bending strength perpendicular to the minimal (Imin) and maximal (Imax) axes of the bone cross sections were statistically increased in GFC rats, as well as the torsional strength along the bone (Fig. 2N-P). Cumulatively, our findings imply that gut microbiota is capable of inducing an acute response resulting in bone mass increase and enhancement in bone mechanical properties.

### Induction of periosteal bone apposition by gut microbiota accounts for increased cortical bone mass

Micro CT measurements of cortical bone at tibia midshaft showed significant bone mass accumulation in GFC rats and implied that periosteal bone apposition was responsible for this increase. However, this analysis was conducted on a thin 0.5 mm tomographic segment of tibia. In order to validate this result on a broader scale and to get an insight into the nature of bone accumulation on a cellular level, we conducted histologic analysis of approximately 12 mm longitudinal sections of tibia diaphysis. Microscopic examinations of these specimens clearly demonstrated noticeable differences between GF and GFC rats in the distribution of circumferential lamellar bone. Measurements of areas occupied by lamellar bone on the periosteal surfaces showed an enlargement of this compartment by 5-fold (Fig. 3A and 3C) with mean thickness increased over 2-fold in GFC versus GF rats (Fig. 3B-C). On the endosteal surfaces, area and mean thickness were quite the same in both animals (Fig. 3A-C). These results indicate that indeed accelerated periosteal bone apposition is likely responsible for increased cortical bone mass in GFC rats. Histologic specimens used for these measurements were cut longitudinally and parallel to anterior-posterior axis of tibia, therefore only anterior and posterior surfaces could be measured in these specimens. To assess the distribution of change in thickness around the tibia shaft, we analyzed 4 tomographic slices spaced evenly along the length and location comparable to histologic specimens. Visualizations of local thicknesses were obtained in these slices using conventional sphere method of distance transformation available in our image analysis software package. Interestingly, the increase in bone thickness in GFC animals was localized to the anteromedial section of the cortex and spanning approximately 1/3 of the bone circumference (Fig. 3D). This section of the tibia receives most of the strain (28). Assuming that GF and GFC animals experience a comparable bone adaptation to habitual mechanical loading, the observed difference in local bone thickness suggests that microbiota accelerates osteogenesis around sites subjected to increased mechanical stimulation.

**Figure 3.**
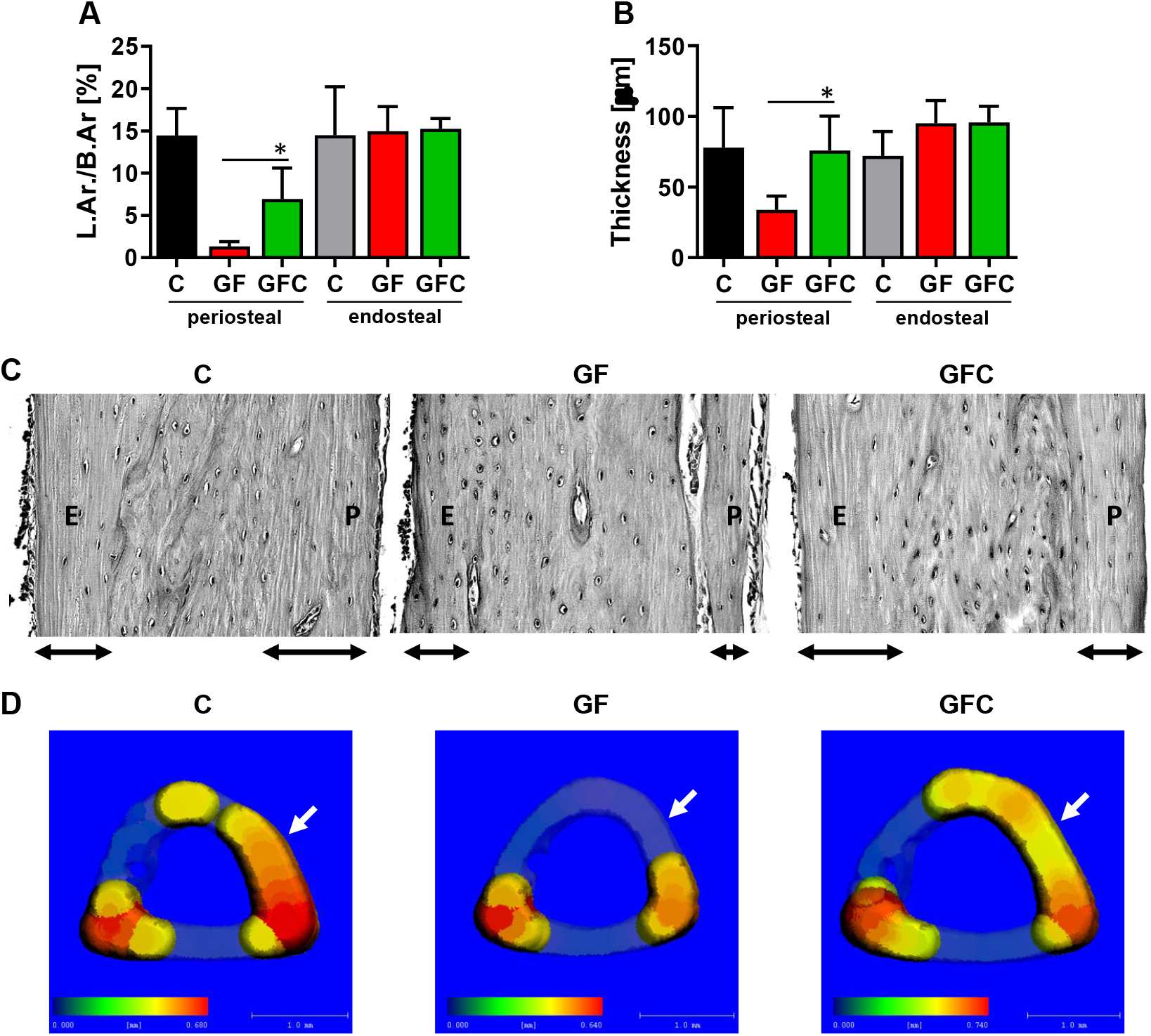
Circumferential lamellar bone apposition in tibia diaphysis of C, GF and GFC rats. A. Percent of periosteal bone and endosteal bone occupied by lamellar bone in analyzed rats. L.Ar./B.Ar. – lamellar bone area as a fraction of either periosteal or endosteal bone area. B. Thickness of lamellar bone in periosteal or endosteal bone. C. Images of representative histologic specimens of tibia cortical bone used for measurements presented in A and B. Two-headed horizontal arrows indicate the extent of lamellar bone. P - periosteal side, E – endosteal side. D. Representative mCT renderings of tibia midshaft cross sections show distribution of bone thickness over 0.44 mm thickness threshold. Blue color in scale bar indicates 0.0 mm thickness, red color indicates maximum thickness for given specimen. Transparent bone sections show thickness below 0.44 mm, white arrows point towards anteromedial bone surface. *: p<0.05

### Microbiota stimulate growth plate enlargement, chondrocyte maturation and longitudinal bone growth

In the course of characterizing a skeletal response to the reconstitution of the holobiome, we measured differences in tibia length between GF and GFC rats and assessed morphology of the epiphyseal plate in proximal tibia. Mean tibia length was significantly increased by 1.6 mm in GFC rats over 10 day of conventionalization period with individual lengths ranging from 35.9 mm to 38.7 mm, while in GF rats the length range was from 34.9 mm to 36.8 mm (Fig. 4A). Reported longitudinal growth rate in Sprague-Dawley rats at 10 weeks of age is 97 + 7 μm per day (29). This comparison shows that tibia longitudinal growth rate was greatly accelerated in GFC rats, which were gaining 160 μm of length per day over an apparently lower growth rate in GF rats. Histologic examinations of decalcified sections of proximal tibia revealed that mean epiphyseal plate thickness was significantly increased in GFC animals by 29% (Fig. 4B) and correlated with dramatic changes in morphology of the plate growth zones (Fig. 4C). Cumulative thickness of the proliferation and prehypertrophic zones in GFC animals was increased by 25%, while hypertrophic zone thickness was increased by 37%, as compared to GF animals (Fig. 4B). Also, a comparison of the epiphyseal plate cellular morphology of C, GF, and GFC animals, as shown in Fig. 4C, strongly suggests that gut microbiome is directly involved in the regulation of longitudinal bone growth by acting on chondrocyte progression and maturation in GFC rats.

**Figure 4.**
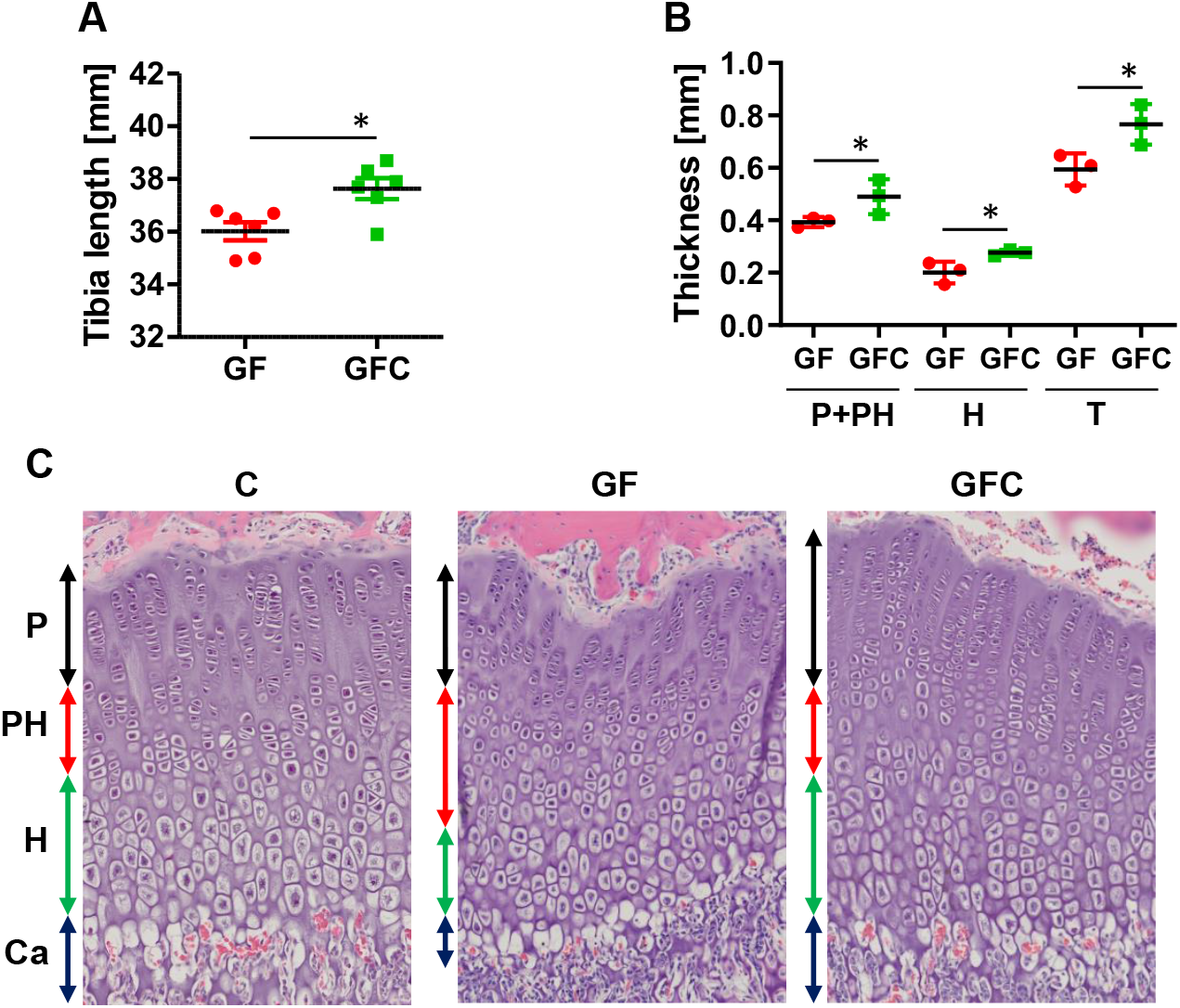
Microbiota-induced growth plate enlargement and improved chondrocyte maturation and calcification promote longitudinal bone growth. A. Total tibia length. B. Growth plate thickness. P+PH – thickness of combined proliferative and prehypertrophic zone; H - hypertrophic zone thickness; T – total thickness. C. Representative histological images of growth plates. Vertical arrows indicate growth plate zones: proliferative (P, black), prehypertrophic (PH, red), hypertrophic (H, green), and calcification (Ca, black). *: p<0.05

### Genes directly involved in bone formation and resorption are not underlying contributors to bone growth facilitated by microbiota

Next, we sought to identify putative effector genes responsible for the observed acceleration of bone mass accumulation in GFC rats by the analysis of gene expression using RNA isolated from bone marrow and bone tissue from tibia diaphysis (Supplementary Fig. 1-2). Distal-less homeobox 5 (*Dlx5*), osterix (*Osx*), osteocalcin (*Bglap*), Collagen 1a1 (*Col1a1*), and sclerostin (*Sost*) were reporters of osteoblast/osteocyte activities and bone formation, whereas periostin (*Postn*) was used as marker of periosteal bone apposition, while receptor activator of nuclear factor kappa-B ligand (*Rankl*) and osteoprotegerin (*Opg*) were reporters of the extent of osteoclastogenesis. We also measured expression of peroxisome proliferator-activated receptor gamma (*Pparg*), adiponectin (*Adipoq*), and fatty acid binding protein 4 (*Fabp4*) to assess bone-related activities of bone marrow adipocytes. Surprisingly, we did not detect any statistically significant differences in the expression of the analyzed genes in both bone tissue and bone marrow in GF and GFC animals (Supplementary Fig. 1-2). At this point, we hypothesized that genes directly involved in bone formation and resorption are less likely to be direct acceptors of regulatory signals resulting from bacterial colonization of the gut, leading us to shift our focus onto systemic regulatory factors.

### Introduction of gut microbiota into germ-free rat affects bone marrow cellular environment

Bone marrow adipose tissue (BMAT) is an essential component of the marrow niche supporting osteogenesis, osteoclastogenesis, and hematopoiesis. BMAT constitutes of adipocytes with different functions depending on their origin and skeletal localization, and represents dynamic tissue rapidly responding to environmental and physiologic cues (30-32). Interestingly, little is known on the effect of microbiota on the BMAT function and the way the holobiont is involved in differentiation and regulation of the marrow adipocyte function. In this study, histologic evaluation of longitudinal sections of tibia diaphysis revealed that reconstitution of holobiont in GF rats increased a density of marrow adipocytes by 39% (Fig. 5A). This difference, although substantial, was not statistically significant. Detailed analysis of adipocyte size distribution showed that this expansion was accompanied with a sequential shift in adipocyte size. As compared to GF rats, there was a significant increase in the count of small in size cells (170-380 μm^2^), which constituted approximately 50% of all adipocytes present in the GFC bone marrow while maintaining histology of adipocytes dispersed in marrow (Fig. 5B). There was no significant difference in the number of large (660-1500 μm^2^) and medium (310-660 μm^2^) sized adipocytes between groups. The reduction in adipocyte size may indicate a shift in adipocyte metabolism toward increased expenditure of stored energy, in the form of released free fatty acids (33,34), in support of a rapid bone accrual.

**Figure 5.**
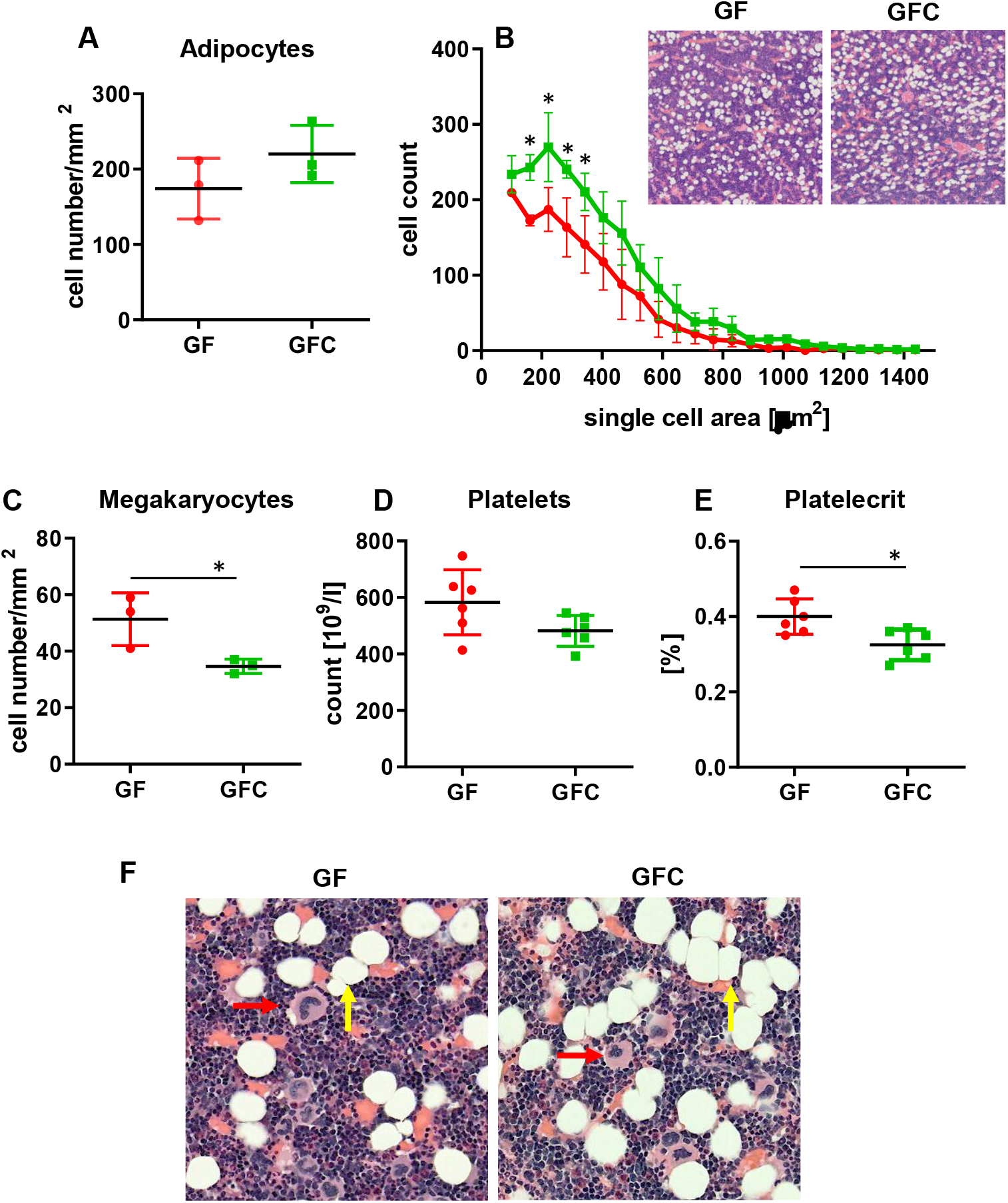
The effect of microbiota on bone marrow adiposity and megakaryocytes. A. Adipocyte cell number per square millimeter of bone marrow in tibia diaphysis. B. Quantitative distribution of adipocyte size in GF (red circles) and GFC (green squares) animals. Inlet represents sections of bone marrow histologic specimens stained with H&E (magnification 10×). C. Number of megakaryocytes per square millimeter of bone marrow in tibia diaphysis. D. Number of platelets in peripheral blood. E. Platelecrit – volume occupied by platelets in peripheral blood. F. Representative histological specimens stained with H&E of marrow in tibia diaphysis showing adipocytes (yellow arrows) and megakaryocytes (red arrows) (magnification 20×). *: p<0.05

Besides a support for bone accrual, it has been shown recently that a direct transfer of fatty acids from marrow adipocytes supports megakaryocyte maturation (35). Considering the shift in adipocyte size, we compared densities of megakaryocyte population with respect to the microbiota status. Surprisingly, the expansion of small in size adipocytes in response to new microbiota seen in GFC rats correlated negatively with the number of mature megakaryocytes (Fig. 5C and 5F). Decrease in the megakaryocyte density in the GFC bone marrow apparently resulted in tendency to a drop in platelet count (Fig. 5D) and significant decrease in plateletcrit in peripheral blood (Fig. 5E). We speculate that this phenomenon may not be associated with adipocytes or reflect a change in adipocyte function, but perhaps toward supporting bone formation at the expense of megakaryocytes maturation. Nevertheless, these results are clearly indicating that gut microbiota shape bone marrow environment supporting adipogenesis and hematopoiesis.

### Microbiota promote host bone growth possibly by the short chain fatty acid butyrate as a metabolite signal for IGF-1 expression

To determine whether any of the microbial-generated metabolites in systemic circulation of the host were different between GF and GFC rats, we conducted a targeted metabolomics analysis of their cecal content. All three short-chain fatty acids (SCFAs), acetate, propionate and butyrate were significantly higher in GFC compared to GF rats (Fig. 6A). Importantly, a dramatic 160-fold increase was observed in the concentration of butyrate in GFC animals compared to GF rats (Fig. 6A). Butyrate is a known metabolite capable of signaling to regulate the IGF-1 anabolic effect on bone (12,36). The *Igf-1* transcript is primarily expressed in liver and IGF-1 biosynthesis and secretion are controlled by growth hormone (GH) and IGF-binding proteins (37,38). *Igf-1* is also locally expressed in bone by osteoblasts and marrow mesenchymal stem cells (39,40). The level of *Igf-1* expression was significantly increased by 3-fold in the liver of GFC rats, whereas in GFC bone and bone marrow expression remained on the same level as in GF rats (Fig. 6B). Interestingly, low IGF-1 protein level in the serum of GF animals was restored to normal, when compared to C rats, after microbiota reconstitution in GFC rats (Fig. 6C). This observation suggests that reconstitution of microbiota results in the *de novo* supply of SCFA including butyrate, which signals *via* modulation of *Igf-1* expression in the liver and restores the threshold of the circulating IGF-1 hormone to the level required for efficient bone formation (36).

**Figure 6.**
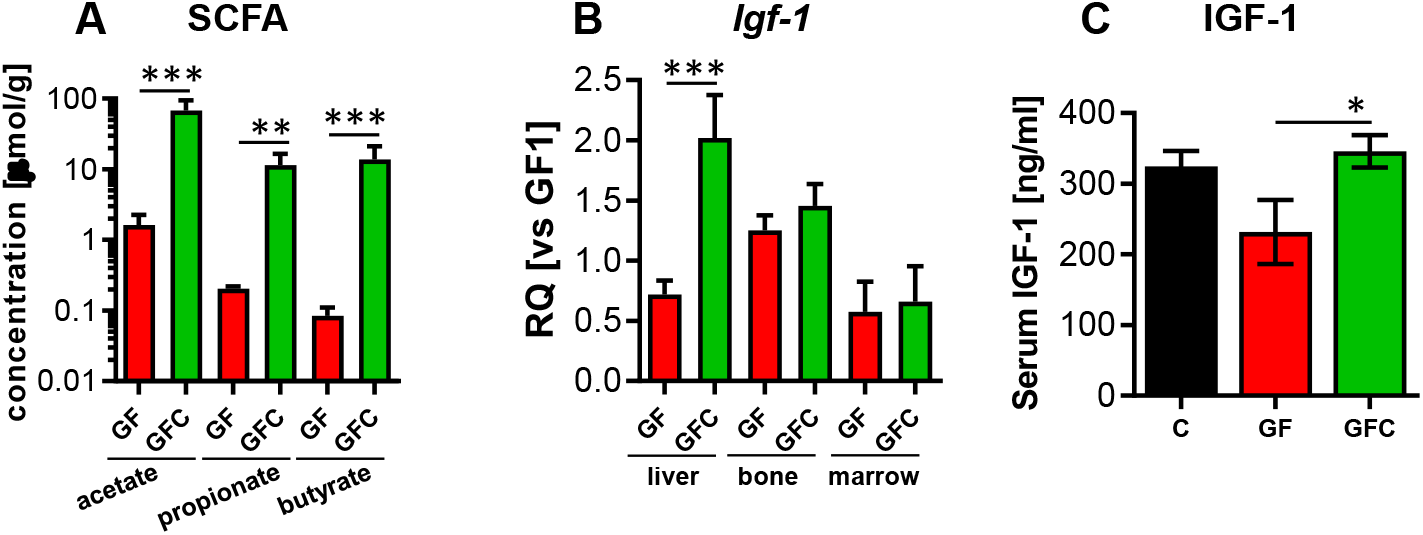
The effect of microbiota on cecal concentration of SCFA and IGF-1 expression. A. Cecal concentrations of acetate, propionate, and butyrate in GF and GFC rats. Note the log 10 scale of Y axis. B. *Igf-1* mRNA expression in liver, bone, and bone marrow in GF and GFC rats. Relative quantity (RQ) was calculated relative to the highest value in GF group (GF1). C. Serum levels of IGF-1 protein in C, GF, and GFC rats. *: p<0.05, **: p<0.01, ***: p<0.001

## Discussion

Recent advances in research addressing systemic host-gut microbiota interactions show that bone is a significant component of the holobiome being strongly tied to the status of gut microbiota. A number of studies showed a positive influence of gut microbiota on bone in animal models of skeletal deficiencies and pathologies, as well as in humans with senile osteoporosis (6,41-44). At the same time, multiple studies described malnutrition associated gut microbiota dysbiosis as a plausible factor for stunted growth in juveniles (45), which attests the gut microbiota as a relevant target and potent remedy for bone-related ailments. However, whether gut microbiota itself is a ‘bystander’ or a ‘mediator’ on bone metabolism has been less clear.

In this paper we show, for the first time, that *de novo* introduction of gut microbiota to GF rat, through cohousing with age- and sex-matched conventional rat, can lead to an abrupt increase in bone mass, acceleration of longitudinal bone growth, and increased bone mineralization. Skeletal changes in GFC rats assessed in tibia bone were manifested by increased midshaft cortical thickness, expansion of trabecular bone, lengthening of tibia, and overall increase in bone size. Interestingly, cortical thickness and the ratio of bone area to total cross-sectional area at tibia midshaft (B.Ar/T.Ar) in individual GF rats was positively correlated with their respective body mass, indicating that a fundamental principle of bone functional adaptation to body mass is maintained even in the absence of gut microbiota (46,47). However, these correlations were negative in GFC animals regardless of the significant increase in their body weight, as compared to GF rats. This indicates that processes regulating bone accrual becomes temporarily independent of the mechanostat (47-49) and that additional mechanism(s) take over to accelerate bone growth and restore a balance between body weight and skeleton, which resembles the naturally occurring adolescent spurt growth (50-52).

To our knowledge, the observed phenotypes are unprecedented considering that the changes occurred within a 10-day time window following gut colonization. Specifically, the bone alterations could be associated with the distinctive alterations in β-diversity observed in GFC animals, particularly with regard to the abundance of *Muribaculaceae, Clostridium* spp., and *Suterella* spp., which may have occurred in response to signals originating from the newly established holobiont in combination with self-adjustment of the microbiota populating microbial-free gut environment of the GF rat. Notably, *Clostridium* spp. are known producers of the short fatty acid (SCFA) butyrate in the colon, and *Muribaculaceae* have recently emerged as a bacterial family potentially possessing this capacity (53), while *Sutterella* spp. encompass a number of low-abundance species characterized by high resistance to toxic effects of bile acids (54). Thus, an increase in relative abundance of these species may aid together development of communities supporting a rapid bone growth in a dysbiotic adolescent organism (22,55,56).

Fermentation of complex carbohydrates in the colon by microbiota results in production of SCFA, acetate, propionate and butyrate, which act as regulatory metabolites on a variety of physiologic processes. Among them butyrate is a prominent modulator of bone homeostasis by affecting the process of bone remodeling. This SCFA was found to protect from postmenopausal and inflammatory bone loss by acting as an inhibitor of bone resorption through shifting preosteoclast cellular metabolism towards glycolysis and by blocking their differentiation through inhibiting histone deacetylase activity (57,58). On the other hand, butyrate induces bone anabolic effect in GF mice and is essential for PTH-dependent bone formation through increasing the number of CD8^+^ T_reg_ cells which produce WNT10b (9). Interestingly, in our study we did not detect any difference in the expression of this ligand in GF and GFC bone marrow suggesting the WNT signaling pathway to be less likely involved in this model. However, we observed that the dramatic increase of cecal butyrate concentration in GFC rats was paralleled with increased levels of circulating IGF-1 and enhanced expression of *Igf-1* transcript in the liver, pointing to the somatotropic axis as one of the targets involved in bone enhancement. It has been shown that butyrate increases intracellular calcium levels and boosts the release of growth hormone (GH), which controls IGF-1 production from pituitary (10). However, significant increase in the expression level of liver *Igf-1* observed in GFC rats may also occur independently of GH and may result from a direct effect of butyrate on hepatic IGF-1 production (12). Interestingly, the *Igf-1* mRNA expression in bone marrow and in bone tissue were on the same level in GF and GFC rats strengthening the possibility that the circulating IGF-1 likely contributed to the accelerated bone formation in GFC animals (59,60).

Skeletal changes in GFC rats can be attributed to the endocrine activity of circulating IGF-1 (28,61,62). Analysis of cortical bone accrual revealed that bone gain in GFC animals was localized to the periosteal surface and formation of circumferential lamellar bone with no change to the endosteum, which is consistent with the overall endocrine effect of IGF-1 (63-65). Similarly, reconstitution of gut microbiota had profound effect on the maturation of chondrocytes in the growth plate and was manifested by expansion of proliferative and hypertrophic zones and overall thickening of the epiphysis. This in turn accelerated longitudinal growth, as differentiation of proliferating chondrocytes into hypertrophic chondrocytes is crucial for bone lengthening (66). Again, maturation of epiphyseal chondrocytes is under the control of IGF-1 (67). It remains to be established whether these effects observed in GFC rats can be attributed to the GH axes in response to SCFA or are independent of GH and involve either direct effect of butyrate on hepatic IGF-1 production or direct effect of SCFA on periosteum and growth plate.

It is significant to note that by virtue of employing the deconstruction of the holobiont in GF rats and reconstruction of the holobiont in GFC rats, our work here has provided a critical insight into the rapid nature of the physiologic interaction between bone and microbiota which leads to rapid bone accrual. The immediate resumption of bone growth following the introduction of microbiota into GF rats suggests that the community of the gut microbial species can act as a vital organ with respect to bone growth, rather than being a passive entity, which produces a variety of regulatory metabolites that are utilized by the host at will. It would appear from our study that in GF animals all the factors required for bone growth are set and waiting on the starting line for the final command. Thus, our work defines microbiota as an ‘environmental factor’ to consider in targeting adolescent bone disorders and clinical management of age-related bone loss.

## Acknowledgements

This work was supported by grants from the NIH to BJ and MVK (HL1430820 and CA219144, respectively), American Diabetes Association Innovative Basic Science Award (1-19-IBS-029) to BLC, and grant from the Crohn’s and Colitis Foundation to PS (Ref.# 522820)

## Authors’ roles

BJ, BLC, MV-K, PJC contributed to experimental design, data analysis, and data interpretation; PJC, BSY, MV-K, RMG, SC, XC, BM, AA, JY, SB conducted the experiments, PJC wrote the manuscript; BLC, MV-K, BJ, PJC, RMG edited the manuscript and take responsibility for the integrity of the data analysis.

