## Supplemental for "Reconstitution of the host holobiont in germ-free rats acutely increases bone growth and affects marrow cellular content"

Supplementary Table 1. Primer sequences for quantitative PCR

| Gene | NCBI Ref. Seq. | Forward Primer (5'>3') | Reverse primer (5'>3') | Amplicon (bp) | Tm (Cº) F / R |
|---|---|---|---|---|---|
| *Adipoq* | NM_144744.3 | CAAGGCCGTTCTCTTCACCT | CCCCATACACTTGGAGCCAG | 116 | 64.0 / 64.1 |
| *Col1a1* | NM_053304.1 | TGGCAACCTCAAGAAGTCCC | ACAAGCGTGCTGTAGGTGAA | 93 | 64.1 / 64.1 |
| *Dlx5* | NM_012943.1 | AAGGCTTATGCGGACTACGG | GCTCGGCCACTTCTTTCTCT | 106 | 63.8 / 64.0 |
| *Fabp4* | NM_053365.1 | AGAAGTGGGAGTTGGCTTCG | ACTCTCTGACCGGATGACGA | 103 | 64.0 / 64.2 |
| *Igf1* | NM_001082477.2 | TGGTGGACGCTCTTCAGTTC | CTCATCCACAATGCCCGTCT | 110 | 63.9 / 64.2 |
| *Bglap* | NM_013414.1 | TGCATTCTGCCTCTCTGACC | GGTAGCGCCGGAGTCTATTC | 113 | 63.9 / 63.6 |
| *Tnfrsf11b* | NM_012870.2 | TGTGAAAGCAGTGTGCAACG | CCAGGCAAGCTCTCCATCAA | 83 | 63.8 / 64.2 |
| *Sp7* | NM_001037632.1 | GGCTGAGGAAGAAGCCCATT | AAGTGGGCTTTCAGATGCGA | 85 | 64.3 / 64.3 |
| *Postn* | NM_001108550.1 | GAAGACCACACAGGGAAGCA | AACGTGAATGACGCCGTTTG | 115 | 64.0 / 63.8 |
| *Pparg* | NM_013124.3 | TCCCGTTCACAAGAGCTGAC | ATAATAAGGCGGGGACGCAG | 107 | 63.9 / 64.0 |
| *Tnfsf11* | NM_057149.1 | TCCTGTACTTTCGAGCGCAG | GTCGAGTCCTGCAAACCTGT | 109 | 63.9 / 64.2 |
| *Sost* | NM_030584.1 | GTACATGCAGCCTTCGTTGC | GGAGGCTCTGGGTACTCTCT | 101 | 63.8 / 64.0 |
| *Wnt10b* | NM_001108111.1 | GAACTGCTCGGCACTAGAGG | AAGGAGAACGCACTCTCACG | 87 | 63.8 / 63.8 |
| *Wnt16* | NM_001109223.1 | ATGAACTGAGTAGCGGCACC | AACACTCGGTCATGTTGCCT | 111 | 64.1 / 64.2 |

Supplementary Table 2. Trabecular bone morphometry in L4 vertebrae

| Animal | TV | BV | BV/TV | Conn.D. | SMI | Tb.N | Tb.Th | Tb.Sp |
|---|---|---|---|---|---|---|---|---|
| GF1 | 18.5336 | 4.1993 | 0.2266 | 71.6535 | 1.7080 | 3.3134 | 0.0847 | 0.3043 |
| GF2 | 17.1157 | 3.3681 | 0.1968 | 61.4056 | 1.9806 | 3.4914 | 0.0787 | 0.2845 |
| GF3 | 18.5268 | 4.0692 | 0.2196 | 79.4792 | 1.5962 | 3.3293 | 0.0792 | 0.3030 |
| GF4 | 20.3986 | 5.1171 | 0.2509 | 88.2903 | 1.2875 | 3.6679 | 0.0811 | 0.2708 |
| GF5 | 15.2616 | 2.9726 | 0.1948 | 70.4711 | 1.8732 | 3.2672 | 0.0774 | 0.3101 |
| GF6 | 16.0442 | 2.9747 | 0.1854 | 66.0987 | 1.9406 | 3.1826 | 0.0761 | 0.3174 |
| **Mean** | **17.6468** | **3.7835** | **0.2124** | **72.8997** | **1.7310** | **3.3753** | **0.0795** | **0.2984** |
| GFC1 | 20.4157 | 4.1111 | 0.2014 | 63.7010 | 1.7580 | 3.1026 | 0.0820 | 0.3232 |
| GFC2 | 21.1751 | 5.1403 | 0.2428 | 83.5178 | 1.3744 | 3.2723 | 0.0837 | 0.3107 |
| GFC3 | 21.3683 | 5.0757 | 0.2375 | 74.3157 | 1.4478 | 3.4275 | 0.0836 | 0.2917 |
| GFC4 | 18.7845 | 4.1512 | 0.2210 | 80.3456 | 1.5221 | 3.2769 | 0.0786 | 0.3070 |
| GFC5 | 19.1017 | 4.1312 | 0.2163 | 70.0461 | 1.4813 | 3.1187 | 0.0812 | 0.3231 |
| GFC6 | 18.2254 | 3.8565 | 0.2116 | 71.5761 | 1.6505 | 3.4540 | 0.0782 | 0.2886 |
| **Mean** | **19.8451** | **4.4110** | **0.2218** | **73.9171** | **1.5390** | **3.2753** | **0.0812** | **0.3074** |
| **T-test** | **0.041** | 0.157 | 0.448 | 0.840 | 0.145 | 0.311 | 0.311 | 0.356 |

Supplementary Table 3.  Tibia midshaft cortical bone morphometry

| **Animal** | **T. Ar.** | **B.Ar.** | **M.Ar.** | **Ct.Th.** | **BMD** | **TMD** |
|---|---|---|---|---|---|---|
| GF1 | 5.1708 | 3.2765 | 1.8943 | 0.4640 | 606.57 | 935.32 |
| GF2 | 5.0096 | 3.2426 | 1.7670 | 0.4220 | 605.84 | 914.34 |
| GF3 | 5.0779 | 3.1683 | 1.9096 | 0.4360 | 594.29 | 927.17 |
| GF4 | 5.4558 | 3.1963 | 2.2595 | 0.4270 | 560.39 | 933.95 |
| GF5 | 4.5712 | 2.6374 | 1.9338 | 0.4060 | 531.16 | 907.10 |
| GF6 | 4.4148 | 2.5955 | 1.8193 | 0.4130 | 552.69 | 914.16 |
| **Mean** | **4.9500** | **3.0194** | **1.9306** | **0.4280** | **575.16** | **922.00** |
| GFC1 | 5.7308 | 3.5176 | 2.2132 | 0.4630 | 585.59 | 930.83 |
| GFC2 | 5.3006 | 3.4303 | 1.8703 | 0.4990 | 631.13 | 953.83 |
| GFC3 | 5.8677 | 3.5989 | 2.2688 | 0.4740 | 593.65 | 946.41 |
| GFC4 | 5.2546 | 3.4098 | 1.8448 | 0.4790 | 623.15 | 941.00 |
| GFC5 | 5.5763 | 3.6413 | 1.9350 | 0.4960 | 630.94 | 945.77 |
| GFC6 | 5.7061 | 3.3526 | 2.3535 | 0.4390 | 563.14 | 934.13 |
| **Mean** | **5.5727** | **3.4917** | **2.0809** | **0.4750** | **604.60** | **941.99** |
| **T-test** | **0.008** | **0.006** | 0.221 | **0.004** | 0.118 | **0.007** |

Supplementary Figure 1. Analysis of gene expression in bone tissue

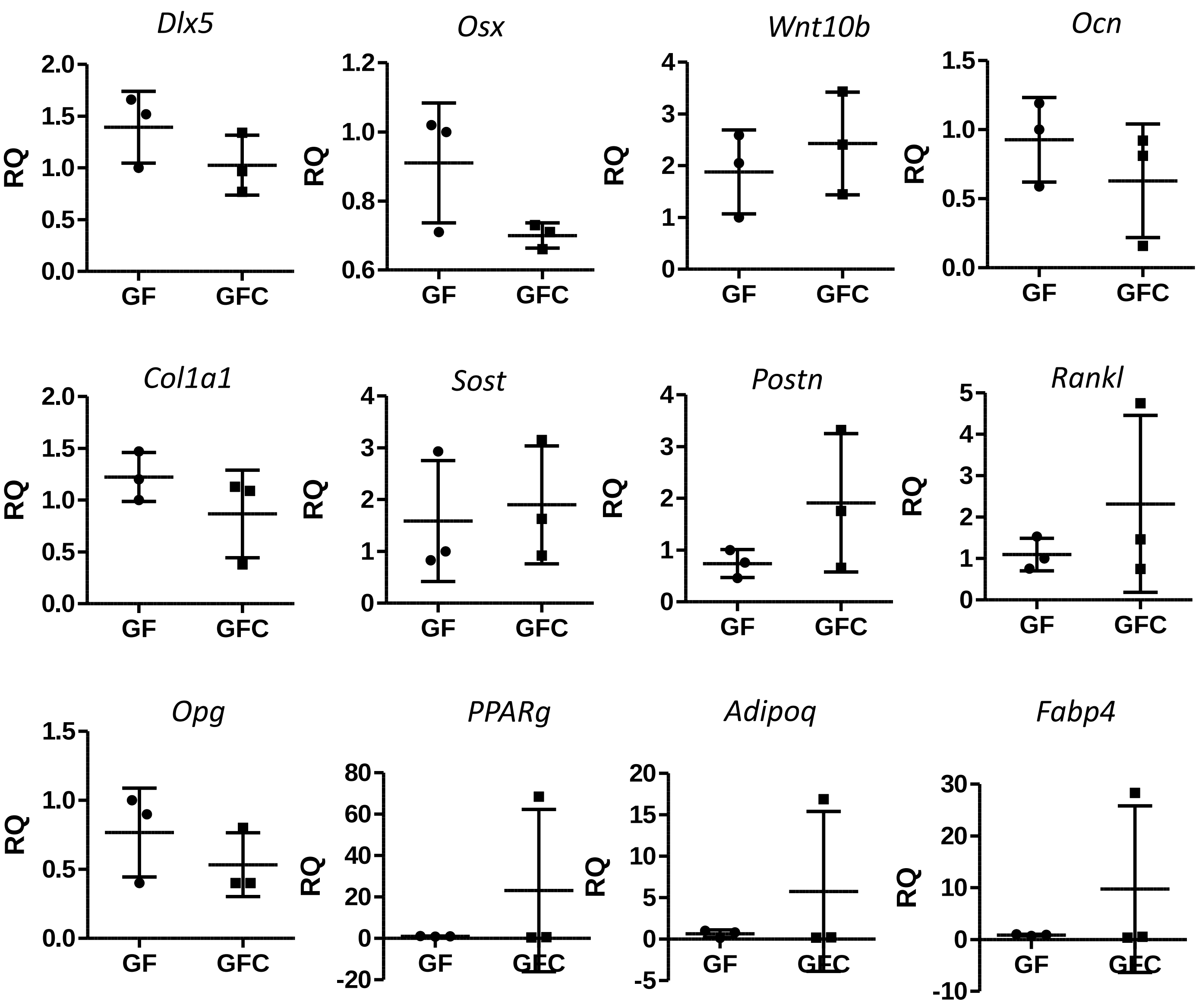

Supplementary Figure 2. Analysis of gene expression in bone marrow

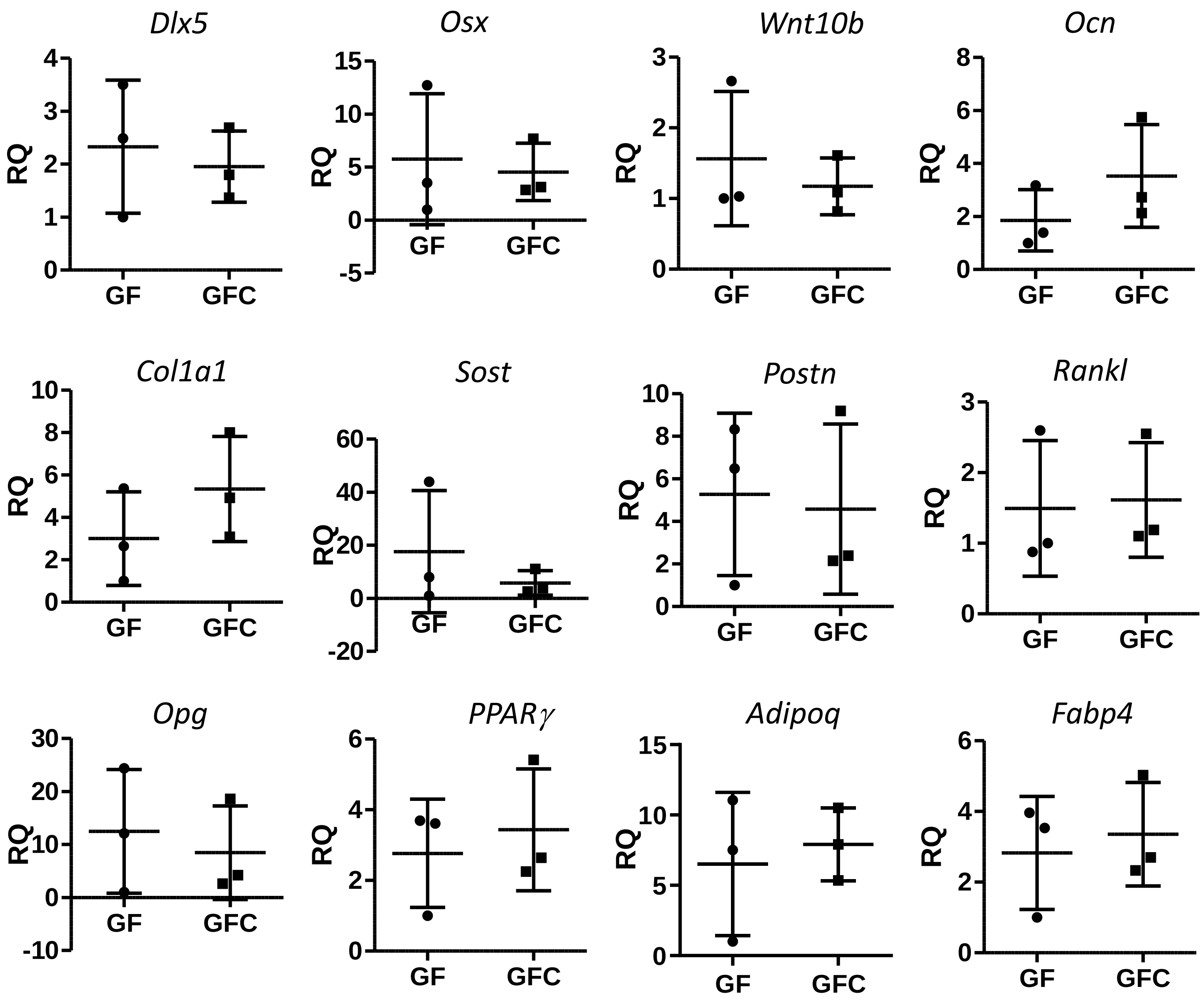
